# Differential evolution of cooperative traits in aggregative multicellular bacterium *Myxococcus xanthus* driven by varied population bottleneck sizes

**DOI:** 10.1101/2023.09.20.558552

**Authors:** Jyotsna Kalathera, Vishwa Patel, Samay Pande

## Abstract

Repeated population bottlenecks influence the evolution and maintenance of cooperation^1,2^. However, it remains unclear whether bottlenecks select all cooperative traits expressed by an organism or only a subset of them. *Myxococcus xanthus*, a social bacterium, displays multiple cooperative traits, including growth, predation, sporulation in multicellular fruiting bodies, and germination ^3–6^. Using laboratory evolution experiment, we investigated the effect of repeated stringent versus relaxed population bottlenecks on the evolution of these four cooperative traits when they were all under selection. We found that only fruiting body formation and growth were positively selected under the stringent regimen, while the other two traits were negatively selected. The pattern was reversed in the relaxed regimen. Additionally, the relaxed regimen led to a significant increase in fitness when competed against ancestors across the entire lifecycle, whereas the stringent treatment did not change competitive fitness. Genomic analysis revealed that mutations in σ^54^ interacting protein and DNA-binding response regulator protein are linked with the changes observed in stringent and relaxed regimens respectively. Further, similar trade-offs are also seen among natural populations of *M. xanthus*. Overall, we demonstrate that different bottleneck sizes drive the evolution of lifecycles in distinct manners, driven by trade-offs between cooperative life history traits.

## Introduction

Examples of cooperative microbial interactions in which cells help their clonemates are abound ^7–9^. However, explaining the evolution and stability of such interactions is challenging ^10,11^. A significant fraction of cooperative interactions are driven by diffusible “public good” molecules that are susceptible to exploitation by non-producers ^12–15^, as they benefit from availability of freely available public goods produced by cooperators in the environment without contributing resources towards their production ^11,16–19^.

Population bottlenecks can stabilise cooperating interactions. In lab evolution experiments, repeated stringent bottleneck have been shown to enhance biofilm formation^1^ and multicellular development in both prokaryotic and eukaryotic microbes ^2,20^. This phenomenon is attributed to the increased kinship between individual ^2,21–26^. We observed that for most studies that demonstrate the effects of population bottlenecks on the evolution and maintenance of cooperation, the model organism could potentially express multiple social traits ^1,2,12,20,27^. However, the focus of investigation has generally been limited to only one or a minority of the subset of these social traits. Since expression of multiple social traits is common ^28–30^, it is crucial to investigate whether stringent bottlenecks positively influence all or only a subset of cooperative traits that a microbe can express.

For microbes that express multiple cooperative traits as part of their lifecycle, we hypothesised that population bottlenecks will probably positively affect the evolution of only some but not all social traits. Population bottleneck events often lead to a reduction in population diversity, resulting in increased kinship among surviving individuals ^20,31^. If a variant that efficiently expresses a cooperative trait survives such bottlenecks ^31,32^, it is likely to become fixed in the population. In contrast, relaxed bottleneck events, due to clonal interference, favour strategies where variants can reap maximum benefits without investing resources ^33,34^. This might create an opportunity for the evolution of cheaters that exploit cooperators expressing costly and essential social traits, as reduced kinship allows for the persistence of such exploitative behaviour ^26^. Conversely, we hypothesise that since the efficiency of expressing less essential cooperative traits might have less impact on evolutionary success, repeated bottleneck events are likely to lead to a decline in the performance of these traits. However, in populations experiencing weaker bottleneck events, less essential traits may be maintained due to limited benefits for cheaters. By investigating the effects of population bottlenecks on the evolution of cooperative traits in a single organism, we aim to shed light on the differential selection pressures acting on various social traits and their implications for the overall biology of the organisms.

To test this, we used *Myxococcus xanthus* as a model organism to study the effect of repeated population bottlenecks on four distinct social traits that were under selection during a lab evolution experiment (supplementary figure 1). *M. xanthus* is a gram-negative soil bacterium with a complex lifecycle. Multiple stages of *M. xanthus* lifecycle were previously shown to be cooperative. These include sporulation ^35^, predation ^4^, germination ^6^ and growth ^3^. We performed lab evolution experiment under two distinct bottleneck regimens. During these experiments all four cooperative traits mentioned above (growth, multicellular spore filled fruiting body formation, germination, and predation) were under selection.

We demonstrate that the essential cooperative traits in the context of the design of the evolution experiment (i.e., sporulation and growth) are selected when populations experience stringent bottlenecks. By exploring the influence of population bottlenecks and importance of a given social trait in the biology of organisms, our study not only provides insights into the evolutionary dynamics of cooperative behaviours but also sheds light on the effect of bottlenecks on the evolution of lifecycles involving multiple cooperative traits.

## Results

### Sporulation and growth are maintained or positively selected for in stringent population bottleneck regimen

To understand the effect of population bottlenecks, we propagated *M. xanthus* populations in conditions in which four distinct social traits (namely growth, sporulation, germination, and predation) were under selection (Supplementary figure 1). During the lab evolution experiment *M. xanthus* populations were propagated for 10 cycles each of which consisted of growth, followed by starvation induced development and sporulation, followed by germination and predation on *E. coli* lawns. Between each of 10 transfer cycles, either 1 % (stringent regimen) or 15 % (relaxed regimen) of the *M. xanthus* population was transferred from predation condition to liquid growth condition (Supplementary figure 1). After 10 cycles of transfer evolved populations and clones were analysed to test our hypothesis that not all but only a subset of cooperative traits will be positively selected under stringent bottleneck treatment.

First, we tested the effect of evolutionary regimen on starvation induced sporulation. Starving *M. xanthus* cells initiates an aggregative developmental process that culminates in the formation of multicellular fruiting bodies filled with spores. Previous studies have shown that population bottlenecks influence the evolution of aggregative behaviours in both *Dictyostellium discodieum*, *M. xanthus* and fungi ^2,20,36^. Consequently, we hypothesized that populations subjected to stringent population bottlenecks would exhibit improved aggregative development compared to populations subjected to relaxed population bottlenecks. To test this, we inoculated both the ancestral isolate and evolved populations onto starvation (TPM) agar plates. In such conditions *M. xanthus* populations develop spore-filled multicellular fruiting bodies, allowing for visual monitoring. This qualitative assessment indicated that all replicate populations from the stringent population bottleneck regimen displayed enhanced fruiting body formation (supplementary figure 2a) compared to populations from the relaxed regimen. Given the identical fruiting body phenotypes observed across replicate populations, we selected population D15 (from the 15 % regimen) and population D1 (from the 1 % regimen) as representative populations for further investigations.

Evolved populations often contain multiple genotypes, and interactions among these genotypes can influence the overall phenotype of the populations. Hence, we checked whether distinct clones isolated from the representative populations exhibit similar fruiting body formation abilities as seen at the population level. For this, we randomly selected three clones from population D1 (from the 1 % regimen) and three clones from population D15 (from the 15 % regimen). A comparison of the developmental phenotypes among these clones revealed similarities to the phenotypes observed in their respective populations of origin (supplementary figure 2b).

Further quantitative analysis of the representative isolates revealed that, on average, clones from the 1 % regimen exhibited a 2.6 log-fold increase in spore production compared to the clones from the relaxed treatment (Figure 1a, independent-sample t test, *p-value* < 0.0009^(FDR^ ^corrected)^). The higher productivity of clones from stringent treatment was not due to increased sporulation efficiency compared to the ancestors, but rather a result of reduced spore productivity in the relaxed regimen clones (Figure 1a). These findings suggest that fruiting body development is negatively selected in the relaxed population-bottleneck regimen, while repeated stringent population-bottleneck events lead to the maintenance of sociality during sporulation.

**Figure 1:**
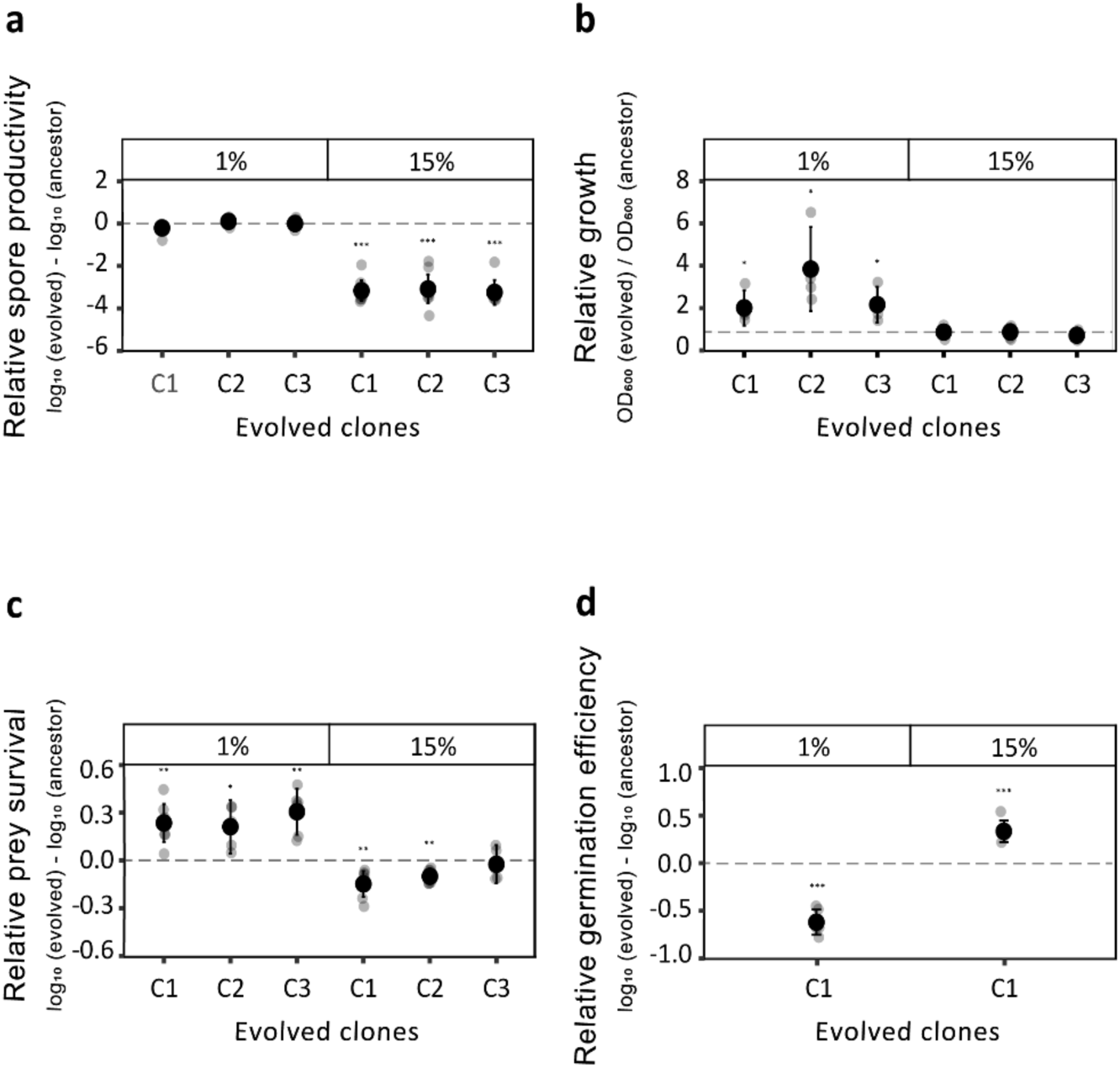
Stringent population bottleneck selects for faster growth and efficient sporulation, whereas relaxed population bottleneck selects for improved predation and germination-efficiency. Spore productivity, growth, predatory performance, and germination efficiency of three representative clones from 1 % and 15 % treatment were measured and plotted relative to the performance of their respective ancestors. (a) Data plotted is the spore productivity of evolved isolates relative to the ancestor (FDR corrected one-sample t test for differences relative to zero, *p* *** < 0.001, n = 7). (b) Clones from stringent population-bottleneck treatment grow more in 48 h relative to the ancestors. Data plotted is O.D. _600 nm_ of evolved isolates relative to the ancestral variant (FDR corrected one-sample t test for differences relative to zero, *p* * < 0.05, n = 3). (c) Growth of prey bacterium *E. coli* was used as a measure of predation efficiency. Data shown is the growth (CFU / mL) of *E. coli* in the presence of evolved clones relative to the ancestors (FDR corrected one-sample t test for differences relative to zero, *p*\*\*\* < 0.001, *p*\*\* < 0.01, *p* * < 0.05, n = 7). (d) Percentage of spores germinated in 4h is used as the measure of germination efficiency. Data represents germination efficiency of evolved clones relative to that of the ancestors (FDR corrected one-sample t test for differences relative to zero, *p* *** < 0.001, n = 6).

Like sporulation, the growth of *M. xanthus* is also a density-dependent trait influenced by the availability of digested proteins and digestive enzymes^3^ which act as public goods. In our evolution experiment, populations underwent growth in a liquid CTT medium before being transferred to the development plate for sporulation. Based on this, we hypothesized that the population from the 1 % regimen, which exhibited improved sporulation efficiency, would also demonstrate a superior growth phenotype. Indeed, clones derived from the 1 % regimen exhibited a 2.5-4.5 -fold higher productivity after 48 hours of growth in a liquid medium compared to clones from the 15 % treatment (independent-sample t test growth of clones from 1 % regimen and 15 % regimen, *p-value* < 0.002^(FDR^ ^corrected)^). Furthermore, the growth of the clones from the 1 % regimen was significantly higher than that of their ancestors, while the growth of clones from the 15 % regimen remained similar to that of their ancestors (Figure 1b). Overall, these experiments provide evidence that populations evolved under stringent conditions outperform their counterparts from the relaxed regimen in terms of both growth and sporulation.

During the evolution experiment, the transfer of *M. xanthus* spores onto *E. coli* lawns subjected the evolving populations to selection pressures for predation and germination on *E. coli*. Predation by *M. xanthus* involves the use of contact-dependent and independent molecules such as antibiotics and digestive enzymes to kill and digest prey cells ^37–40^. These killing molecules, along with the extracellularly digested dead prey, contribute to the public good. Hence, we hypothesized that the stringent regimen, which promotes higher kinship, would result in improved performance during predation and germination, similar to development and growth.

To assess the predatory performance of evolved and ancestral *M. xanthus* isolates, *M. xanthus* cells were co-cultured with *E. coli* under conditions similar to the evolution experiment. These experiments revealed that the *M. xanthus* isolates evolved in the 15 % treatment exhibited higher predation efficiency compared to the isolates evolved in the 1 % regimen (2.3-fold lower CFU / mL of *E. coli* in the presence of *M. xanthus* from the 15 % treatment compared to the clones from the 1 % treatment, independent-sample t test, *p-value* < 0.014^(FDR^ ^corrected)^). Furthermore, comparison between the ancestors and the evolved clones showed that relaxed population bottlenecks selected for increased predatory performance (on average 1.3-fold lower CFU / mL relative to the ancestors, one-sample t test, *p-value* < 0.006^(FDR^ ^corrected)^ or had no effect on the predatory performance (one out of three isolates tested)), while stringent population bottlenecks selected for either decreased performance (two out of three isolates) relative to the ancestors (on average 1.8-fold higher CFU / mL of *E. coli* in the presence of *M. xanthus* relative to the ancestors, one-sample t test, *p-value* < 0.03^(FDR^ ^corrected)^) (Figure 1c).

Germination of *M. xanthus* spores is a density-dependent cooperative trait driven by diffusible public goods ^20^. Due to the complexity and time-sensitive nature of measuring germination efficiency, we focused on studying one representative isolate from each treatment for this analysis. The estimation of germination efficiency in the evolved and ancestral isolates showed that the clones from the 15 % regimen exhibited significantly higher germination rates (on average 100 % spores germinated within 4 hours) compared to the ones from the 1 % regimen (on average 0 % spores germinated) (Figure 1d, independent-sample t test *p-value* < 1.041e-06). These results further demonstrate that when multiple cooperative traits are under selection, only a few are positively selected or maintained in a stringent bottleneck regimen.

The differential performance of clones from the 1 % and 15 % regimens in growth, sporulation, germination, and predation could be attributed to the maintenance of cooperative traits or the evolution of variants that excel in these specific traits without relying on social interactions. To assess whether these traits involve social interactions, we utilized density dependence as a valuable tool. Cooperative behaviours often display density-dependent effects, with the benefits or expression of the trait increases with higher population densities ^41,42^. By examining how the traits respond to varying population densities, we can infer the presence of cooperation within the bacterial population. Thus, to demonstrate the cooperative nature of growth and sporulation in the clones from 1 % regimen, as well as the cooperative predation in the 15 % regimen clones, we tested for density dependent performance of the evolved isolates for these respective traits. Due to logistical constraints, we were unable to measure the density dependence of germination efficiency (see methods). These experiments revealed strong positive density dependence in growth (ancestor : *R sq.* = 0.6822, *p-value* < 2.2e-16; 15 % : *R sq.* = 0.5484, *p-value* = 7.621e-07; 1 % : *R sq.* = 0.5305, *p-value* = 1.445e-06) and sporulation efficiency (ancestor : *R sq.* = 0.7536, *p-value* < 2.2e-16; 15 % : *R sq.* = 0.03303, *p-value* = 0.1952; 1 % : *R sq.* = 0.5522, *p-value* = 9.051e-11) for the 1 % regimen clones. Similarly, the 15 % regimen clones exhibited positive density dependence in predation efficiency (ancestor: *R sq.* =-0.4765, *p-value* = 5.563e-08; 15 %: *R sq.* = -0.5488, *p-value* = 1.127e-09; 1 %: *R sq.* = -0.0521, *p-value* = 0.8636). Moreover, as expected, the ancestral variant displayed positive density dependence in each of the four traits (Figure 2).

**Figure 2:**
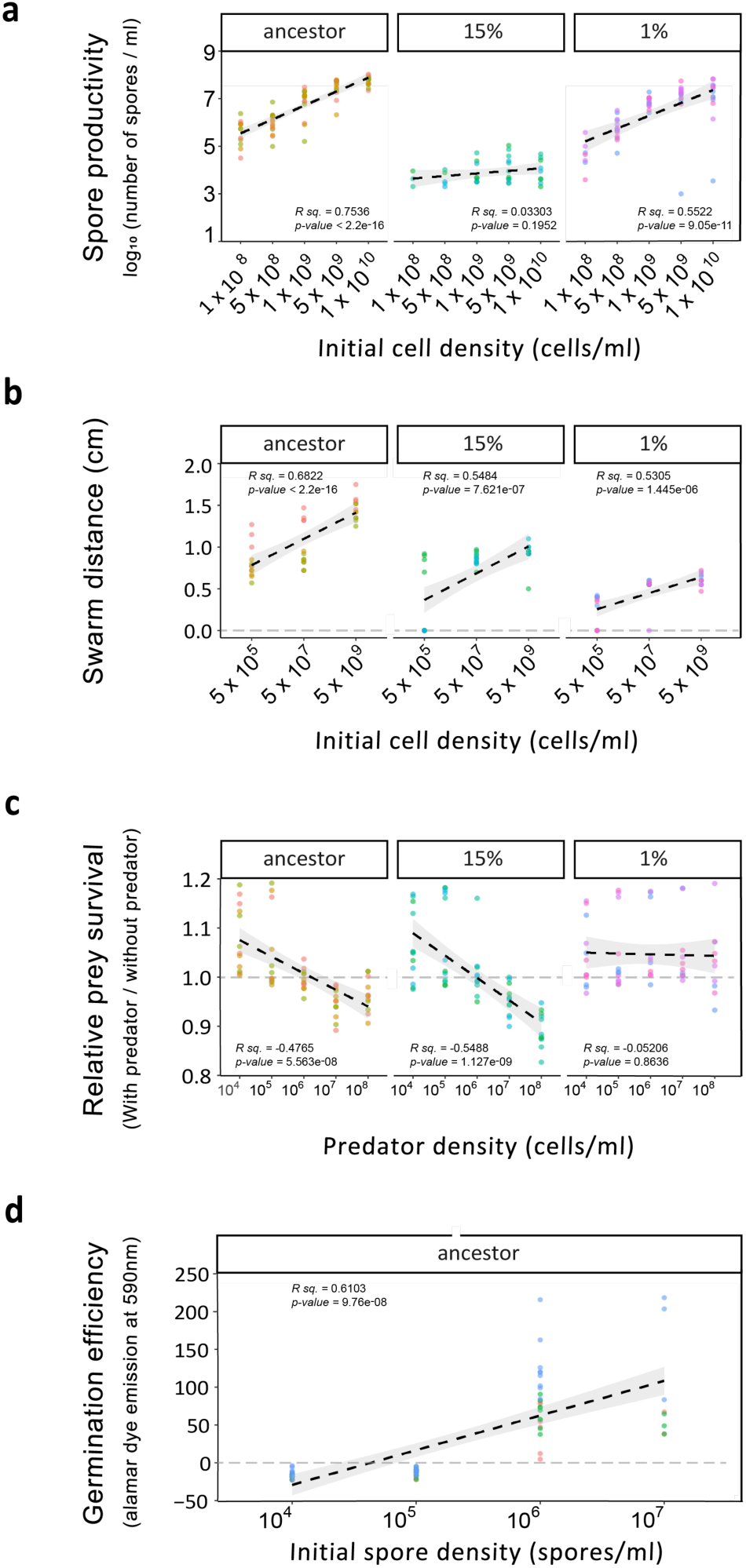
Social traits in *M. xanthus* are positive density dependent. Dots represent efficiency values of respective trait measured for three different colonies each of ancestors, 15 % evolved and 1 % evolved populations. Dashed lines are fitted linear regression and shaded area are 95 % confidence interval a) Sporulation productivity for the ancestral strain GV1, 15 % evolved and 1 % evolved clones on starvation media (TPM-hard agar) is shown. Slope is significantly positive for ancestors and isolates derived from 1 % selection regimen. (n = 4) (ancestor : *R sq.* = 0.7536, *p-value* < 2.2e-16; 15 % : *R sq.* = 0.03303, *p-value* = 0.1952; 1 % : *R sq.* = 0.5522, *p-value* = 9.051e-11) b) Swarm expansion (growth) for the ancestral strain GV1, 15 % evolved and 1 % evolved clones on agar bed (CTT-hard agar) is shown. Slope is significantly positive for ancestors and isolates derived from 1 % and 15 % selection regimen (n = 4). The swarm expansion is density-dependent in ancestor, 15 % and 1 % regimen. (ancestor: *R sq.* = 0.6822, *p-value* < 2.2e-16; 15 % : *R sq.* = 0.5484, *p-value* = 7.621e-07; 1 % : *R sq.* = 0.5305, *p-value* = 1.445e-06) C) Predation efficiency of ancestral *M. xanthus* clones, and evolved clones from 15 % and 1 % treatments was measured as the growth of *E. coli* when it was cocultured with *M. xanthus* at four different densities. Negative slope of the regression line indicates increasing predation efficiency with increasing density (n = 4). Significant negative slope in 15 % show that predation by these isolates is a density dependent social trait. (n = 4) (ancestor : *R sq.* =-0.4765, *p-value* = 5.563e-08; 15 % : *R sq.* = -0.5488, *p-value* = 1.127e-09; 1 % : *R sq.* = -0.0521, *p-value* = 0.8636) d) Germination efficiency of ancestral *M. xanthus* isolates was measured as a function of increasing spore density. Exit from dormancy and metabolic activity of the spores is measured as emission at 590 nm when spores are inoculated in nutrient rich suspension with alamar blue (n = 4) (ancestor: *R sq.* = 0.6013, *p-value* = 9.76e-08). For all traits analysed ancestor clones exhibit positive cell density dependence.

Taken together, distinct cooperative traits were either maintained or improved in the two regimens. Importantly, the traits enriched in one treatment were selected against in the other. Thus, demonstrating that the size of population bottleneck can determine which cooperative traits are selected.

### A relaxed population bottleneck selects for higher competitive fitness across the lifecycle, but a stringent bottleneck does not

It is commonly observed in evolution experiments that the loss of cooperative traits often gives rise to cheaters who exploit the cooperators. In line with this understanding, we proposed that the loss of cooperation in the evolved variants would coincide with their ability to cheat the ancestral strain specifically in the focal trait. To support this notion, we conducted a competition experiment between clones that had evolved under the 15 % regimen with reduced sporulation efficiency and the ancestral strain during starvation-induced sporulation. The results of these experiments revealed a significant 2.3-fold fitness advantage for the clones evolved in the 15 % regimen over the ancestors when mixed in a 1:1 proportion (Figure 3a, one-sample t test, *p-value* < 1.819e-08^(FDR^ ^corrected]^). These results demonstrate the cheating behaviour of the evolved variants towards the ancestors. These findings strongly suggest that the selection of such variants may be driven by the benefits associated with cheating.

**Figure 3:**
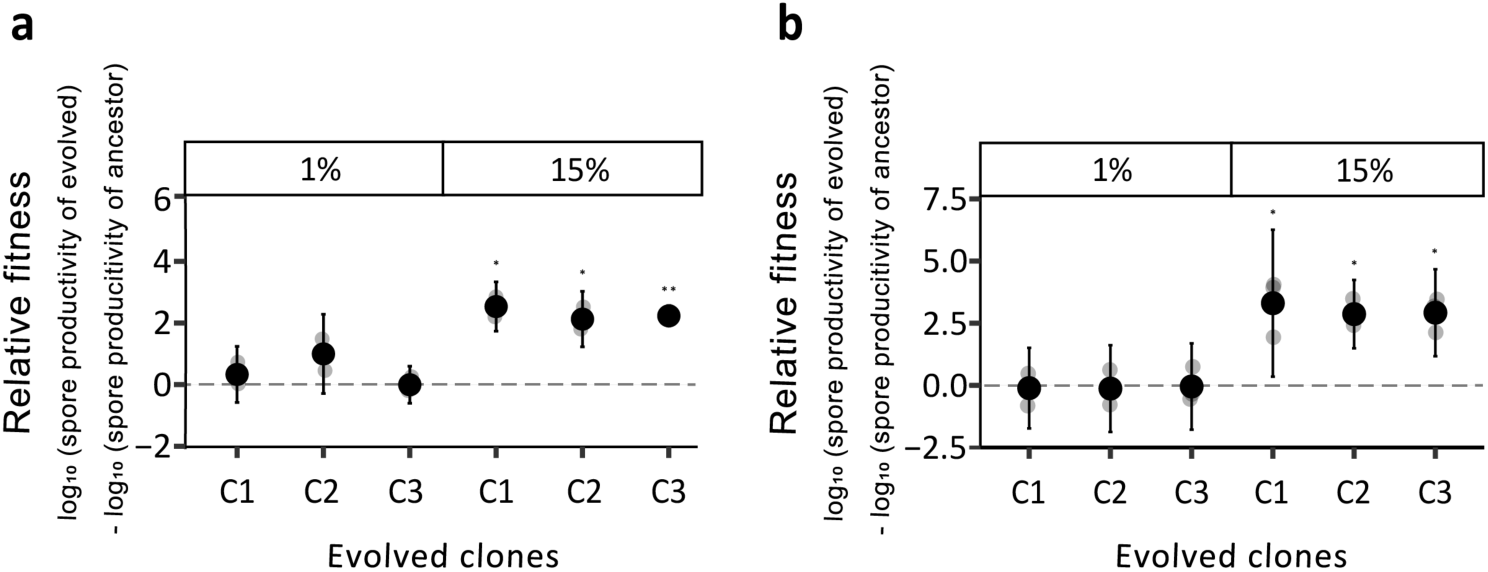
Evolved strains are either equally fit or are fitter than the ancestor when competed across either fruiting body developmental cycle or entire lifecycle. Representative clones from stringent (1 %) and relaxed (15 %) regimen were mixed with the ancestors in 1:1 proportion and relative fitness of evolved clones against the common ancestor was assessed. a) Clones from 1 % treatment were equally fit as their ancestors, whereas clones from 15 % regimen showed higher relative fitness during competition during one developmental cycle (FDR corrected one-sample t test for differences relative to zero, p ** < 0.01, *p* * < 0.05, n = 3). b) Similar to developmental competition, during competition across two rounds of life cycle, clones from 1 % treatment were equally fit as their ancestors, whereas clones from 15 % regimen showed higher relative fitness (FDR corrected one-sample t test for differences relative to zero, *p* * < 0.05, n = 3).

**Figure 4:**
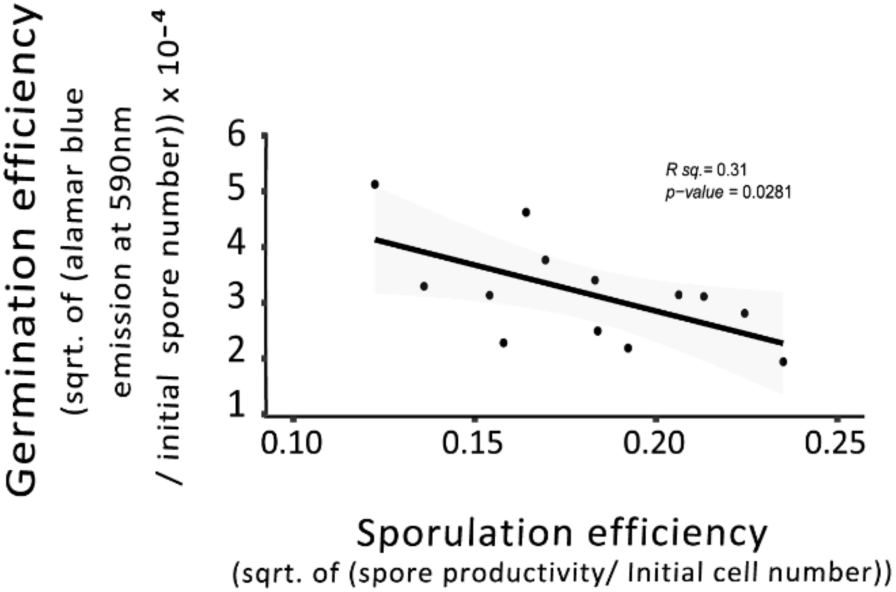
Spore productivity and germination efficiency are negatively correlated in natural *M. xanthus* populations. Spore productivity of 13 randomly chosen soil derived *M. xanthus* isolates was measured in the conditions similar to the ones used during the evolution experiment and correlated with their germination efficiency. Dashed line indicates fitted linear regression (*R. sq.* =0.31, and *p* < 0.03, n = 4).

We observed that the evolved isolates from stringent and relaxed regimen evolved to perform better at two distinct sets of social traits. These findings suggested that the populations from the two regimens have evolved to survive using two distinct strategies. However, better or worse performance at individual trait may or may not result in competitive advantage for the evolved variants against their ancestors when they compete with each other across lifecycle that involves all traits examined in our experiments. Hence, the evolved isolates were mixed with their ancestors in 1:1 proportion and propagated across the lifecycles twice before their numbers were estimated to count competitive fitness. Interestingly we observed that the isolates from 15 % regimen outcompeted their ancestors repeatedly across replicates (3.66-log-fold higher productivity relative to ancestors one-sample t test *p-value* < 5.38e-06 ^[FDR^ ^corrected]^). Whereas the ones from 1 % regimen on average showed similar fitness to their ancestors (One-sample t. test *p-value* > 0.05). Together, isolates from 15 % regimen seems to have evolved higher degrees of adaptive fitness over their ancestors relative to the ones from 1 % treatment.

### Mutations in regulatory genes are responsible for the changes

Representative clones were sequenced to identify the mutational changes responsible for the observed phenotypes. We found a small number of mutational differences between ancestors and evolved isolates (supplementary table 1). Among the mutational differences identified, only one distinct mutation was present in all isolates of each of the two regimens. Isolates from 1 % regimen had a frameshift mutation in a sigma 54 interacting transcriptional regulation MXAN_4899. This protein was previously shown to be responsible for the regulation of secondary metabolites and hence the predatory performance of *M. xanthus* ^43^. Since this protein in its native form helps in the production of secondary metabolites, it is understandable that a frameshift mutation (potential loss of function) might be responsible for the reduced predatory performance of the isolates from 1 % regimen. Though, why this mutation might affect the growth of the isolates positively is not clear, we suspect that cost saved by not producing the secondary metabolites involved in predation might be responsible for the increased growth of these mutants.

A similar analysis of clones from 15 % regimen revealed that all three isolates acquired a missense mutation. This mutation was responsible for the change in threonine to methionine at the seventy-seventh position of the DNA binding response regulator MXAN_1093. It was previously demonstrated that MXAN_1093 plays a role in the starvation-induced development of *M. xanthus* fruiting bodies. Thus, it is highly likely that the mutation identified in this gene has a negative effect on its functionality and responsible for the low spore productivity of isolates from 15 % regimen. Together, genomic analysis has revealed two mutations that are very likely to be the causative agents for the changes observed in the evolved isolates.

### Life-history traits of natural isolates are negatively correlated, indicating the presence of trade-offs in nature

Demonstrating the significance of laboratory observations in natural settings has been emphasised numerous times. Therefore, we tested whether similar to the lab-evolved populations of *M. xanthus*, natural populations also demonstrate trade-offs between sporulation (i.e., development) and germination. We decided to use these two traits because they showed the highest levels of divergence between populations evolved in 15 % and 1 % regimen (figure 1a, figure 1d). To perform this analysis, we randomly picked thirteen natural isolates of *M. xanthus*. Estimates of germination efficiency using alamar dye, and starvation-induced sporulation efficiency revealed a significant negative correlation between the two traits (R. sq. = 0.31, *p-value* < 0.03). These findings imply that the trade-off we observed between spore productivity and germination efficiency may be a general constraint on *M. xanthus* social evolution in nature that drives the evolution of lifecycles of *M. xanthus* in natural populations.

## Discussion

Explaining the evolution and maintenance of cooperative interactions poses a challenge. Previous studies have shown that population bottlenecks play a crucial role in stabilizing cooperative behaviour, primarily by increasing kinship among individuals ^1,2,20,44^. However, most of these studies have focused on examining a single cooperative traits ^1,2,20^. Considering that many microbes possess multiple social traits ^28–30^, we sought to investigate whether stringent population bottlenecks would select for all cooperative traits or only a few. The results presented demonstrate that when stringent bottlenecks are repeatedly applied, only a select few cooperative traits are favoured. Interestingly, the cooperative traits of sporulation and growth, which are either maintained or positively selected under stringent conditions, do not exhibit improvement in the relaxed regimen. Conversely, in the relaxed condition, germination and predation are favoured.

In line with numerous other microbial species, *M. xanthus* exhibits the ability to express multiple social traits, such as swarming, growth, sporulation, germination, and predation. These traits are predominantly driven by diffusible substances ^3,6,45,46^ and, in some cases, additional contact-dependent interactions ^5,39^. Each of these social traits can be studied under distinct environmental conditions, enabling us to replicate the life cycle of *M. xanthus* in a laboratory evolution experiment. In this experiment, *M. xanthus* populations were grown in a nutrient-rich environment, transferred to starvation agar for development, and then transferred to growing prey lawns for germination and predation. Thus, sporulation, germination, predation, and growth were all subjected to selection.

While all four social traits were under selection, growth and sporulation experienced more stringent selection pressures compared to germination and predation. This is because, after being spotted on starvation agar, all cells that failed to transition into spores were eliminated before transfer to the prey lawn. Hence, the ability to sporulate was essential for survival. Once inoculated on the prey lawn, spores could afford to germinate slowly and possess less efficient predation mechanisms since, after four days of coincubation, the cultures were transferred to a nutrient-rich liquid medium. Superior growth in the liquid medium ensured that individuals would be transferred to the starvation agar plate for the next round of the life cycle. Therefore, performance in terms of growth and sporulation was more critical than achieving higher efficiency in germination or predation.

Genetic drift strongly influences adaptation when populations experience stringent bottlenecks. Under such conditions, the first beneficial mutations that arise by chance face minimal clonal competition and are more likely to fixate, especially when selection pressure is high ^32,33,47,48^. In the evolution experiment conducted in this study, populations underwent stringent bottlenecks just before being transferred to a growth medium. Thus, any variant that exhibited superior fitness during the growth phase in the liquid medium would increase in frequency and survive if it also possessed the ability to sporulate. Our results strongly support this evolutionary dynamic as the driving force behind the evolution of variants that perform better during liquid growth and sporulation induced by starvation under stringent treatment compared to the relaxed treatment. As expected from this hypothesis, the variant enriched in our experiment demonstrated both strong growth capabilities and maintained similar levels of sporulation efficiency as its ancestor, resulting in its fixation.

In contrast, under relaxed bottleneck treatment, higher levels of genetic diversity and clonal interference enable the selection of the fittest variants. Our results demonstrate that efficient sporulation is not the optimal competitive strategy. Instead, the ability to sporulate better in the presence of an efficient sporulator is favoured. This finding aligns with previous research, suggesting that variants capable of exploiting cooperative traits outcompete individuals investing resources in those traits. For these exploitative variants to succeed, it is not essential for them to increase in frequency during growth. Rather, their ability to exploit sporulating strains during starvation is sufficient for their transfer to the next environment, such as *E. coli* lawns for germination and predation. Our results further suggest that clonal interference among exploiters of developmental cooperators on *E. coli* leads to the emergence and fixation of efficient germinators and predators.

Taken together, our study demonstrates that when multiple social traits are under selection, stringent population bottlenecks select for the maintenance of the essential cooperative traits and negatively impact the performance of non-essential social traits. In contrast, relaxed population bottlenecks select for reduced performance and the evolution of exploitation in essential cooperative traits, while favouring increased efficiency in non-essential social traits. Thus, we provide evidence that stringent bottlenecks do not necessarily result in increased cooperation among individuals and can actually lead to reduced cooperativity. Considering that most microbial species exhibit multiple social traits ^27,49^ these findings hold broad relevance.

Complex lifecycles are common, ranging from aggregative multicellularity seen in *M. xanthus* and *D. discodieum* to lifecycles involving two developmental stages, such as larval development followed by metamorphosis or terminal development ^50,51^. Our observations suggest that population bottlenecks within such lifecycles influence the overall evolution of life history strategies. Therefore, based on this study, it would be intriguing to explore the role of population bottlenecks on the evolution of lifecycle.

## Materials and Methods

### Strains and Culture conditions

*Myxococcus xanthus* bacterial strains used in the study were GV1, and GV2 (Rifampicin resistant version of GV1) obtained from Gregory Velicer, ETH Zurich ^52^. The *Escherichia coli* strain used in the study was laboratory strain MG1655 ^53^. All *M. xanthus* strains were stored in CTT liquid media ^54^ containing 20 % glycerol at - 80 °C. Whereas *E. coli* was stored in LB medium with 20 % glycerol at - 80 °C.

To obtain *M. xanthus* cultures for assays, the strains were inoculated from frozen stocks on CTT hard agar (1.5 % agar, 1 % casitone, pH 7.6) plates ^54^, incubated at 32 °C for three days during which time the swarms of *M. xanthus* appeared on the plates. Edges of the three-day old swarms were inoculated in 8 mL CTT liquid (1 % casitone, pH 7.6) media ^54^ in 50 mL conical flasks, and incubated at 32 °C with constant shaking at 200 rpm till they reached desired O.D. _600 nm_ (0.2 - 0.8).

To obtain spores, cultures grown in CTT liquid medium were centrifuged at 5000 rpm for 20 min at 25 °C and cell pellets were resuspended to 5×10^9^ cells / mL density using TPM buffer (pH 7.6)^45^ and spotted (100 µL, until otherwise specified) on TPM hard agar (1.5 % agar, pH 7.6) plates, followed by incubation at 32 °C for three days. Fruiting bodies were harvested in 1 mL ddH_2_O using sterile scalpel, then heated at 50 °C for two hours, and sonicated for 20 seconds (Amplitude: 25, Pulse: 10 sec ON, 10 sec OFF and 10 sec ON) using Q700 sonicator (Qsonica) with 24 tip horn (part #4579)

*E. coli* cultures were initiated by streaking the glycerol stock on LB agar medium, followed by incubation at 32 °C overnight. A single colony from the LB agar plate was inoculated in 8 mL Luria Broth in 50 mL conical flask and grown overnight until O.D. _600 nm_ 1 - 1.2 and washed once and further adjusted to 0.1 O.D. _600 nm_ using TPM buffer to use in all predation assays.

### Isolation of natural *M. xanthus* isolates

To obtain natural isolates of *M. xanthus*, soil samples were obtained from various locations of Indian Institute of Science (IISc) campus, Bengaluru, India. To collect soil, top cut sterile 10-mL syringes were used. Following the removal of a ∼2mm section of top soil from inside the syringe with a sterile scalpel, the remaining soil was crushed well and covered on top of a selective medium ^55^ (TPM hard agar (1.5 % agar) with 0.5 % casitone, vancomycin (10 µg / mL) (Sigma Aldrich, V2002, CAS number 1404-93-9), nystatin (1 U / mL) (Sigma Aldrich, N4014 50MG, CAS number 1400-6-19), cycloheximide (50 µg / mL) (Sigma Aldrich, 01810-5G, CAS number 66-81-9) and crystal violet (10 µg / mL) (Sigma Aldrich, C0775, CAS number 548-62-9). The soil-covered plates were incubated at 32 °C for over two weeks, until fruiting bodies appeared. Thirteen different fruiting bodies from distinct locations were randomly collected and transferred to separate microcentrifuge tubes with sterile 1 mL ddH_2_0 with a toothpick. These samples were incubated at 50 °C for 2 hours to kill any vegetative cells and enrich only thermoresistant spores. These samples were then sonicated for 20 seconds (Amplitude: 25, Pulse: 10 sec ON, 10 sec OFF and 10 sec ON) to release the spores from fruiting bodies before diluting and plating on CTT-soft agar (0.5 % agar) media. The spores germinated on CTT-soft agar plates over the course of 4 - 5 days were then collected and used for CTT-liquid inoculation, where each colony forming unit on the soft agar plate representing a spore. Individual genotypes were designated as X.Y, where X denotes the fruiting body from which it originated, and Y represents the distinct spore number from that single fruiting body. The cultures were kept at 32 °C and 200 rpm until they reached 0.2 - 0.8 (O.D. _600 nm_) and then frozen in - 80 °C with 20 % glycerol.

### Isolation of *M. xanthus* clones from evolved population of *M. xanthus*

In order to better understand within population dynamics, we isolated clones from evolved populations as follows; The evolved population glycerol stocks were serially diluted and plated on CTT soft agar (0.5 % agar) media in 90 mm petri-plates. After 4 - 5 days of incubation at 32 °C, individual colonies (three each from the ancestor and evolved populations) were picked and used to inoculate 8 ml of CTT-liquid media in a 50 mL conical flask from the plates with more dispersed and diverse colonies. When the grown cultures reached an O.D._600 nm_ of 0.2 - 0.8, they were stocked in 20 % glycerol and stored at - 80 °C. The three clones from each treatment were named C1, C2, and C3.

### Experimental evolution

Conditions and sequence of transfers, and the protocol for the evolution experiment is broadly illustrated in the Supplementary figure 1. To replicate different life stages of *M. xanthus* lifecycle three distinct growth conditions were used. For vegetative growth 50 mL conical flasks with 8 mL CTT liquid medium (1 % casitone, pH 7.6) was used. For development and sporulation TPM hard agar (1.5 % agar) beds were made by pouring 10 mL of TPM hard agar (1.5 % agar) in 60 mm petri dish (Tarsons, ref: 460061). For germination and predation phase, spores of *M. xanthus* were co-inoculated with *E. coli* on 10 mL TPM hard agar (1.5 % agar) medium with 0.025 % glucose in 50 mL conical flasks. Since glucose is the only carbon source, *M. xanthus* relies on its predatory abilities to grow by using growing population of *E. coli* as the sole source of nutrients. Four replicate lines were propagated for each bottleneck condition, and to do so four GV1 *M. xanthus* colonies were used as distinct ancestors for each replicate population.

To initiate evolution experiment, individual *M. xanthus* colony was inoculated in 8 mL CTT liquid medium in 50 mL conical flasks for each replicate population, grown until the O.D. _600 nm_ reached 0.3 - 0.4, and the cell density was adjusted 5×10^9^ cells / mL using TPM buffer after centrifuging and pelleting at 5000 rpm for 20 min at 25 °C. 200 µL of the density adjusted culture was spotted on TPM hard agar (1.5 % agar) plates, and incubated for three days at 32 °C. Three days of incubation was enough for *M. xanthus* cells to aggregated and to form spore-filled multicellular fruiting bodies. After three days, TPM hard agar plates were incubated at 50 °C for two hours, which results in the death of vegetative cells but not the spores. Surviving spores were harvested by scrapping the agar surface using sterile scalpel and resuspended in 1 mL TPM buffer (pH 7.6). Resuspended spore and *E. coli* suspension was inoculated on 10 mL TPM hard agar (1.5 % agar) with 0.025 % glucose in 50 mL conical flasks and spread with the help of 5 - 7 glass beads by shaking for 5 minutes. Flasks were incubated for 4 days at 32 °C. After four days of incubation, the cultures were harvested with the help of 5 - 7 glass beads after adding 4 mL TPM buffer and shaking at 200 rpm for 30 min. Next, according to the size of population bottleneck either 40 µL or 600 µL culture volume was transferred to 8 mL CTT (with gentamycin 50 µg / mL) liquid medium in 50 mL conical flasks, incubated at 200 rpm at 32 °C till the cultures reached the O.D. _600 nm_ of 0.3 to 0.4. *M. xanthus* is naturally gentamycin resistant bacterium whereas the *E. coli* strain used in this study was gentamycin sensitive, hence conditions used here allowed the growth of only *M. xanthus* and not *E. coli*. Thus, the population size of *M. xanthus* was adjusted to be same for both treatment at the end of every lifecycle. CTT grown cultures were centred down after centrifuging for 20 min at 5000 rpm at 25 °C and adjusted to the density of 5 x 10^9^ cells / mL using TPM buffer and 200 µL of the same was spotted on TPM hard agar (1.5 % agar) plates to initiate the next cycle. The experiment was performed for ten cycles. At every alternate cycle, glycerol (20 % glycerol) stocks were made after growth in CTT (with gentamycin 50 µg / mL) liquid medium and before the initiation of next round of lifecycle and stored at - 80 °C for analysis.

### Development assay

Aliquots from glycerol stocks, for both ancestors and evolved clones, were spotted on CTT hard agar (1.5 % agar) plates. After 3 days of incubation at 32 °C the swarm edges were used to inoculate 8 mL CTT liquid medium in 50 mL conical flasks. For evolved populations, the cultures were obtained by inoculating a small aliquot of the freezer stock directly in CTT liquid medium. These cultures were incubated at 32 °C in 200 rpm in 50 mL conical flasks till the O.D. _600 nm_ reached 0.3 - 0.4, and the cultures were centred down at 5000 rpm for 20 min centrifugation at 25 °C and the density was adjusted to 5 x 10^9^ cells / mL using TPM buffer and spotted on TPM hard agar (1.5 % agar) plates (100 µL) for standard development assays. For qualitative analysis of the development proficiency of the evolved populations plates were imaged after three days of incubation at 32 °C.

For quantitative analysis, after incubating for three days of incubation at 32 °C, plates were baked at 50 °C for two hours, and spores were scrapped of using sterile scalpel, resuspended in 1 mL ddH_2_O, and sonicated (Amplitude: 25, Pulse: 10 sec ON, 10 sec OFF and 10 sec ON). Sonicated spore suspensions were serially diluted, 100 µL of ten-fold dilutions were plated onto CTT soft agar, incubated at 32 °C for 7 days, and colonies were counted.

### Predation assay

Aliquots from glycerol stocks, for both ancestors and evolved clones, were spotted on CTT hard agar plates. After 3 days of incubation at 32 °C the swarm edges formed were used to inoculate 8 mL CTT liquid medium in 50 mL conical flasks. *M. xanthus* culture in CTT liquid media was incubated at 32 °C at 200 rpm, till they reached a O.D. _600 nm_ 0.2 – 0.8. Next, *M. xanthus* cells were centred down (5000 rpm for 20 min at 25 °C) and adjusted to 5 x 10^5^ cells / mL using TPM Buffer.

We measured growth of *E. coli* in the presence of *M. xanthus* as a measure of predatory performance. To do so, *E. coli* cultures were revived from glycerol stock on LB agar plate, a single colony was inoculated in 8 mL liquid LB in 50 mL conical flasks for overnight growth, cultures were then washed in TPM buffer, and adjusted to 0.1 O.D._600 nm_.

All predation assays were performed on 10 mL TPM hard agar (1.5 % agar) beds supplemented with 0.025 % glucose in 50 mL conical flasks with 5 - 7 sterile glass beads on the surface of the agar bed. 50 µL of density adjusted culture of *E. coli* (0.1 O.D. _600 nm_) and 50 µL of density adjusted *M. xanthus* (5 x 10^5^ cells / mL) were co-inoculated and spread on the agar bed with sterile glass beads. Similarly, monocultures of *E. coli* were inoculated as a control which gave us the estimate of the growth of *E. coli* in the absence of *M. xanthus*. For this, 50 µL of density adjusted culture of *E. coli* (0.1 O.D. _600 nm_) was inoculated together with 50 µL of TPM buffer on agar beds and spread using 5 - 7 glass beads. Cultures were incubated at 32 °C for 3 days. Following the incubation period, 4 mL TPM buffer was added to the conical flasks and culture beds were washed by shaking the flasks with the help of 5 - 7 glass beads for 30 min at 200 rpm. Viable counts of *E. coli* were determined by dilution plating in LB soft agar (0.5 % agar) plates.

### Spore germination assay (Ancestors and lab-evolved *M. xanthus* isolates)

Each strain used for germination assay had a specific sporulation efficiency as measured using the sporulation assay mentioned above (See supplementary table 2) for sporulation efficiencies. Thus, we predicted the number of spores expected from the size of starting inoculum of each of the strain used in germination assays. Hence to obtain 10^4^ spores / mL dense spore suspensions of ancestral isolate, spores from one starvation plate (100 µL of 5 x 10^9^ cells / mL inoculum per plate) were harvested in 1 mL ddH_2_O and subsequently serially diluted to get the desired initial spore density mentioned. For clone 1 from 15 % regimen, the spores from 10 starvation plates (100 µL of 5 x 10^9^ cells / mL inoculum per plate) were harvested in 1 mL ddH_2_O to obtain the initial spore density of 10^4^ spores / mL. For clone 1 from 1 % regimen, spores from one starvation plate (100 µL of 5 x 10^9^ cells / mL inoculum per plate) were harvested in 1 mL ddH_2_O and subsequently serially diluted to obtain desired starting density of 10^4^ spores / mL. To confirm that the number of expected spores is same as the actual number of spores used for germination assay, spore suspensions were plated after serial dilution on CTT soft agar (0.5 % agar) plates, incubated at 32 °C for 3 - 4 days, and colonies were counted. These counts were also used as the initial spore numbers inoculated in germination assays (T_0_ spore counts).

To analyse the lab-evolved isolates and their ancestors, germination assays were performed using *E. coli* as the only source of nutrients. To do so, *E. coli* cultures were obtained by inoculating a single colony of *E. coli* in LB media, which was incubated at 200 rpm at 32 °C for overnight. Grown cultures of *E. coli* were resuspended to 0.1 O.D. _600 nm_ in TPM buffer after one round of wash with TPM buffer. 300 µL of 0.1 O.D. _600 nm_ adjusted *E. coli* cultures were inoculated with 10^4^ spores in 1.5 mL microcentrifuge tubes and incubated at 32 °C for 4 h. 100 µL of the coculture was transferred to 900 µL ddH_2_O, heated for 2 hours at 50 °C and sonicated twice for 10 s (Amplitude: 25, Pulse: 10 sec ON, 10 sec OFF and 10 sec ON). 100 µL of sonicated culture was dilution plated onto CTT soft agar (0.5 % agar) plates. Heat treatment and sonication kills spores that have germinated into cells and have lost resistance to sonication as well as heat. Hence the colonies appearing on CTT soft agar (0.5 % agar) plates represent the number of spores that did not germinate in 4 h (T4). Difference between initial spore count and final spore count was used to calculate germination efficiency.

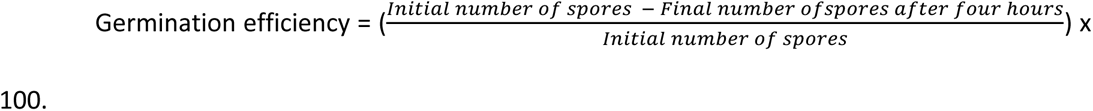

### Spore germination assay (Natural isolates of *M. xanthus*)

Since *M. xanthus* (for isolates used in this study) spore germination is rather fast, the germination assays described above are time sensitive and hence are not ideal for high-throughput analysis of multiple strains of *M. xanthus*. Therefore, to analyse the germination efficiencies of 13 natural isolates of *M. xanthus*, we used an alamar dye-based (alamarBlue™ Cell Viability Reagent, cat. No DAL1025) germination assay which works on the principle of colour change of alamar-dye under a reducing environment, indicating germination of spores into metabolically active vegetative cells. To perform germination assays, 270 µL of 10^6^ spores / mL suspensions of spores were incubated with 30 µL alamar blue dye (10x stock concentration) in 96-well micro-titre plate, incubated at 32 °C in the plate reader for 4.5 h, during which fluorescence intensities (excitation at 550 nm and emission at 590 nm) were measured at 590 nm for every 5 min.

### CTT swarming assay

Swarming distance has been used as a proxy of growth in the past in multiple *M. xanthus* studies. Because of the logistical ease of doing these experiments we used swarming distance as a measure of growth in our density experiments. Swarming assays were performed as detailed in previous studies ^56^. In brief, each clone (15 %: C1, C2, C3 and 1 %: C1, C2, C3, ancestor: C1, C2, C3) was revived from glycerol stocks and inoculated in CTT liquid media. These cultures were incubated at 32 °C at 200 rpm, in 50 mL conical flasks till they reached a O.D. _600 nm_ 0.2 – 0.8. The cells were centred at 5000 rpm for 20 minutes at 25 °C and adjusted to different densities such as 5 x 10^5^ cells / mL, 5 x 10^7^ cells / mL and 5 x 10^9^ cells / mL using TPM buffer. For each isolate, 10 µL of the density adjusted cultures were spotted on the centre of 90 mm petri dish containing solidified CTT hard agar. The spotted plates were incubated at 32 °C, edge of the swarm was marked after 1 day of incubation and then again after 5 days of incubation. The average swarm distance across four distinct axes on each plate between day 5 and day 1 were used as a measure of swarming ability of the respective clones.

### Growth curve assay

*M. xanthus* isolates were revived from glycerol stocks and grown in 8 mL CTT liquid medium in 50 mL conical flasks at 32 °C at 200 rpm for until O.D. _600 nm_ 0.2 – 0.8, centrifuged down at 5000 rpm for 20 min at 25 °C, adjusted to the density of 5 x 10^9^ cells / mL using TPM buffer. 10 µL of the density adjusted culture was transferred to 1 mL CTT liquid in a 48 well microtiter plate (Tarsons, Cat. No. 980051), sealed using transparent cover (Bio-rad, MSB1001), and incubated in Tecan plate reader (Infinite M Nano) at 32 °C for 48 h. O.D. _600 nm_ of the *M. xanthus* cultures was measured every 5 min and cultures were shaken for 10 sec between every reading for aeration.

### Lifecycle competition assay

GV2 (rifampicin antibiotic resistant version of GV1), evolved and ancestor clone cultures were revived from glycerol stocks as mentioned above. 0.2 – 0.8 O.D. _600 nm_ cultures of the *M. xanthus* isolates were centred (5000 rpm, 20 min, 25 °C) and the densities were adjusted to 5 x 10^9^ cells / mL using TPM buffer. 100 µL of density adjusted cultures of the evolved isolate was mixed with 100 µL of density adjusted GV2 culture, and 200 µL of the cocultures were spotted on a TPM hard agar (1.5 % agar) plate and was propagated for two cycles of lifecycle competition using the protocols same as the one used for the evolution experiment. However, the cultures were not subjected to any bottleneck event. At the end of third round of development, the fruiting body spots were harvested using sterile scalpel, and resuspended in 1 mL ddH2O, sonicated twice for 10 s (Amplitude: 25, Pulse: 10 sec ON, 10 sec OFF and 10 sec ON), and dilution plated on CTT soft agar (0.5 % agar) with and without rifampicin. GV2 is a rifampicin resistant variant of GV1 and hence colonies on plates with rifampicin reflect the population size of GV2 (ancestor) whereas the plates without rifampicin represent total number of spores (GV2 + evolved isolate).

### Whole genome sequencing of ancestors and evolved clones

The ancestors and evolved clones (from T_10_ cycle) of the experimental evolution were directly inoculated from respective glycerol stocks to 8 mL CTT liquid in 50 mL conical flasks. The 0.4 - 0.8 O.D. _600 nm_ grown cultures were centrifuged at 5000 rpm for 10 minutes at 25 °C and the cell pellets were subjected for genomic DNA isolation workflow using Qiagen’s Genomic DNA extraction buffer kit (cat. no. 19060) and 20 / G genomic tips. The eluted DNA was stored in 30 µL autoclaved Milli-Q water. The quantity of the DNA was initially checked by using Nanodrop and later at the sequencing facility (Macrogen.Inc, South Korea) using Qubit flurometer. The sequencing was performed in Illumina HiSeq4000 system using Truseq nano DNA kit (350) in paired-end mode generating 150 bp read length, and the samples were prepared according to NGS library preparation workflow. Read quality was assessed using FastQC, and before processing the data the Illumina specific adapters or primers were trimmed using Trimmomatic v0.3365 with the following parameters: ILLUMINACLIP:Nextera+TruSeq3-PE2.fa:2:25:10 CROP:124 HEADCROP:5 LEADING:30 TRAILING:28 SLIDINGWINDOW:4:28 MINLEN:77. For mutational calling, the processed reads were then mapped to a modified version of *M. xanthus* DK1622 genome (available in NCBI with refseq: NC_008095^57^) using breseq pipeline from Barrick’s lab ^58^. We used default clonal analyses parameters described in the breseq documentation with bowtie2 ^59^ as the alignment tool against the reference.

### Statistical analysis

R (version 4.3.1) was used to conduct all the data analyses and for making figures. The figure legends include a description of the statistical test. Every experiment was run in three or more independent replicates separated in different time blocks.

## Acknowledgements

The authors thank DST-FIST for support to the Department of Microbiology and Cell Biology, UGC Centre for advanced study, and the DBT-IISc partnership. This study was supported by the funds from India Alliance Intermediate fellowship to SP (IA/I/20/1/504921).

## Author contribution

SP conceived the project. JK, VP, and SP designed experiments. JK, and VP performed experiments. JK analyzed the data. JK wrote the first draft of the manuscript. SP edited the manuscript.

## References

1. Brockhurst, M. A. Population bottlenecks promote cooperation in bacterial biofilms. PloS One 2, e634 (2007).

2. Kuzdzal-Fick, J. J., Fox, S. A., Strassmann, J. E. & Queller, D. C. High relatedness is necessary and sufficient to maintain multicellularity in *Dictyostelium*. Science 334, 1548–1551 (2011).

3. Rosenberg, E., Keller, K. H. & Dworkin, M. Cell density-dependent growth of *Myxococcus xanthus* on casein. J. Bacteriol. 129, 770–777 (1977).

4. Muñoz-Dorado, J., Marcos-Torres, F. J., García-Bravo, E., Moraleda-Muñoz, A. & Pérez, J. Myxobacteria: moving, killing, feeding, and surviving together. Front. Microbiol. 7, 781 (2016).

5. Kaplan, H. B. & Plamann, L. A. *Myxococcus xanthus* cell density-sensing system required for multicellular development. FEMS Microbiol. Lett. 139, 89–95 (1996).

6. Pande, S., Pérez Escriva, P., Yu, Y.-T. N., Sauer, U. & Velicer, G. J. Cooperation and cheating among germinating spores. Curr. Biol. 30, 4745–4752.e4 (2020).

7. Celiker, H. & Gore, J. Cellular cooperation: insights from microbes. Trends Cell Biol. 23, 9–15 (2013).

8. Crespi, B. J. The evolution of social behavior in microorganisms. Trends Ecol. Evol. 16, 178–183 (2001).

9. Stuart A. West, P. Diggle, S., Angus Buckling, Gardner, A. & S. Griffin, A. The social lives of microbes. Annu. Rev. Ecol. Evol. Syst. 38, 53–77 (2007).

10. Sachs, J. L., Mueller, U. G., Wilcox, T. P. & Bull, J. J. The evolution of cooperation. Q. Rev. Biol. 79, 135–160 (2004).

11. Pande, S. et al. Privatization of cooperative benefits stabilizes mutualistic cross-feeding interactions in spatially structured environments. ISME J. 10, 1413–1423 (2016).

12. Butaitė, E., Baumgartner, M., Wyder, S. & Kümmerli, R. Siderophore cheating and cheating resistance shape competition for iron in soil and freshwater *Pseudomonas* communities. Nat. Commun. 8, 414 (2017).

13. Gore, J., Youk, H. & van Oudenaarden, A. Snowdrift game dynamics and facultative cheating in yeast. Nature 459, 253–256 (2009).

14. Kostylev, M. et al. Evolution of the *Pseudomonas aeruginosa* quorum-sensing hierarchy. Proc. Natl. Acad. Sci. U. S. A. 116, 7027–7032 (2019).

15. Kehe, J. et al. Positive interactions are common among culturable bacteria. Sci. Adv. 7, eabi7159 (2021).

16. West, S. A., Griffin, A. S. & Gardner, A. Evolutionary explanations for cooperation. Curr. Biol. 17, R661–672 (2007).

17. Velicer, G. J. Social strife in the microbial world. Trends Microbiol. 11, 330–337 (2003).

18. Velicer, G. J., Kroos, L. & Lenski, R. E. Developmental cheating in the social bacterium *Myxococcus xanthus*. Nature 404, 598–601 (2000).

19. Chen, R., Déziel, E., Groleau, M.-C., Schaefer, A. L. & Greenberg, E. P. Social cheating in a *Pseudomonas aeruginosa* quorum-sensing variant. Proc. Natl. Acad. Sci. U. S. A. 116, 7021–7026 (2019).

20. Amherd, M., Velicer, G. J. & Rendueles, O. Spontaneous nongenetic variation of group size creates cheater-free groups of social microbes. Behav. Ecol. 29, 393–403 (2018).

21. Hamilton, W. D. The genetical evolution of social behaviour. I. J. Theor. Biol. 7, 1–16 (1964).

22. Foster, K. R., Fortunato, A., Strassmann, J. E. & Queller, D. C. The costs and benefits of being a chimera. Proc. Biol. Sci. 269, 2357–2362 (2002).

23. Nowak, M. A. Five rules for the evolution of cooperation. Science 314, 1560–1563 (2006).

24. West, S. A., Griffin, A. S., Gardner, A. & Diggle, S. P. Social evolution theory for microorganisms. Nat. Rev. Microbiol. 4, 597–607 (2006).

25. Belcher, L. J., Dewar, A. E., Ghoul, M. & West, S. A. Kin selection for cooperation in natural bacterial populations. Proc. Natl. Acad. Sci. U. S. A. 119, e2119070119 (2022).

26. Gilbert, O. M., Foster, K. R., Mehdiabadi, N. J., Strassmann, J. E. & Queller, D. C. High relatedness maintains multicellular cooperation in a social amoeba by controlling cheater mutants. Proc. Natl. Acad. Sci. U. S. A. 104, 8913–8917 (2007).

27. Sathe, S. & Nanjundiah, V. Complex interactions underpin social behaviour in *Dictyostelium giganteum*. Behav. Ecol. Sociobiol. 72, 167 (2018).

28. Brown, S. P. & Taylor, P. D. Joint evolution of multiple social traits: a kin selection analysis. Proc. Biol. Sci. 277, 415–422 (2010).

29. Özkaya, Ö., Balbontín, R., Gordo, I. & Xavier, K. B. Cheating on cheaters stabilizes cooperation in *Pseudomonas aeruginosa*. Curr. Biol. 28, 2070–2080.e6 (2018).

30. Ross-Gillespie, A., Dumas, Z. & Kümmerli, R. Evolutionary dynamics of interlinked public goods traits: an experimental study of siderophore production in *Pseudomonas aeruginosa*. J. Evol. Biol. 28, 29–39 (2015).

31. Wein, T. & Dagan, T. The effect of population bottleneck size and selective regime on genetic diversity and evolvability in bacteria. Genome Biol. Evol. 11, 3283–3290 (2019).

32. Wahl, L. M. & Gerrish, P. J. The probability that beneficial mutations are lost in populations with periodic bottlenecks. Evol. Int. J. Org. Evol. 55, 2606–2610 (2001).

33. Szendro, I. G., Franke, J., de Visser, J. A. G. M. & Krug, J. Predictability of evolution depends nonmonotonically on population size. Proc. Natl. Acad. Sci. U. S. A. 110, 571–576 (2013).

34. Gerrish, P. J. & Lenski, R. E. The fate of competing beneficial mutations in an asexual population. Genetica 102–103, 127–144 (1998).

35. Rosenbluh, A., Nir, R., Sahar, E. & Rosenberg, E. Cell-density-dependent lysis and sporulation of *Myxococcus xanthus* in agarose microbeads. J. Bacteriol. 171, 4923–4929 (1989).

36. Bastiaans, E., Debets, A. J. M. & Aanen, D. K. Experimental evolution reveals that high relatedness protects multicellular cooperation from cheaters. Nat. Commun. 7, 11435 (2016).

37. Sudo, S. & Dworkin, M. Bacteriolytic enzymes produced by *Myxococcus xanthus*. J. Bacteriol. 110, 236–245 (1972).

38. Xiao, Y., Wei, X., Ebright, R. & Wall, D. Antibiotic production by myxobacteria plays a role in predation. J. Bacteriol. 193, 4626–4633 (2011).

39. Seef, S. et al. A Tad-like apparatus is required for contact-dependent prey killing in predatory social bacteria. eLife 10, e72409 (2021).

40. Arend, K. I., et al. *Myxococcus xanthus* predation of Gram-positive or Gram-negative bacteria is mediated by different bacteriolytic mechanisms. Appl. Environ. Microbiol. 87, e02382–20 (2021).

41. Darch, S. E., West, S. A., Winzer, K. & Diggle, S. P. Density-dependent fitness benefits in quorum-sensing bacterial populations. Proc. Natl. Acad. Sci. U. S. A. 109, 8259–8263 (2012).

42. Fiegna, F., Pande, S., Peitz, H. & Velicer, G. J. Widespread density dependence of bacterial growth under acid stress. iScience 26, 106952 (2023).

43. Volz, C., Kegler, C. & Müller, R. Enhancer binding proteins act as hetero-oligomers and link secondary metabolite production to myxococcal development, motility, and predation. Chem. Biol. 19, 1447–1459 (2012).

44. Pedroso, M. The impact of population bottlenecks on the social lives of microbes. Biol. Theory 13, 190–198 (2018).

45. Kuspa, A., Kroos, L. & Kaiser, D. Intercellular signalling is required for developmental gene expression in *Myxococcus xanthus*. Dev. Biol. 117, 267–276 (1986).

46. Berleman, J. E. & Kirby, J. R. Deciphering the hunting strategy of a bacterial wolfpack. FEMS Microbiol. Rev. 33, 942–957 (2009).

47. Patwa, Z. & Wahl, L. M. Fixation probability for lytic viruses: the attachment-lysis model. Genetics 180, 459– 470 (2008).

48. Wahl, L. M., Gerrish, P. J. & Saika-Voivod, I. Evaluating the impact of population bottlenecks in experimental evolution. Genetics 162, 961–971 (2002).

49. Jiricny, N. et al. Loss of social behaviours in populations of *Pseudomonas aeruginosa* infecting lungs of patients with cystic fibrosis. PloS One 9, e83124 (2014).

50. Liedtke, H. C., Wiens, J. J. & Gomez-Mestre, I. The evolution of reproductive modes and life cycles in amphibians. Nat. Commun. 13, 7039 (2022).

51. Bonett, R. M. & Blair, A. L. Evidence for complex life cycle constraints on salamander body form diversification. Proc. Natl. Acad. Sci. U. S. A. 114, 9936–9941 (2017).

52. Kaiser, D. Social gliding is correlated with the presence of pili in *Myxococcus xanthus*. Proc. Natl. Acad. Sci. U. S. A. 76, 5952–5956 (1979).

53. Saha, S., Bhat, B., Laloo, J. M. & Pande, S. Community mixing selects for predation resistance in lab-evolved communities of bacterial prey and social predator *Myxococcus xanthus*. biorxiv, DOI:10.1101/2023.03.21.533597 (2023).

54. Hodgkin, J. & Kaiser, D. Cell-to-cell stimulation of movement in nonmotile mutants of *Myxococcus*. Proc. Natl. Acad. Sci. U. S. A. 74, 2938–2942 (1977).

55. Kraemer, S. A. & Velicer, G. J. Endemic social diversity within natural kin groups of a cooperative bacterium. Proc. Natl. Acad. Sci. U. S. A. 108 suppl. 2, 10823–10830 (2011).

56. Mendes-Soares, H. & Velicer, G. J. Decomposing predation: testing for parameters that correlate with predatory performance by a social bacterium. Microb. Ecol. 65, 415–423 (2013).

57. Goldman, B. S. et al. Evolution of sensory complexity recorded in a myxobacterial genome. Proc. Natl. Acad. Sci. U. S. A. 103, 15200–15205 (2006).

58. Deatherage, D. E. & Barrick, J. E. Identification of mutations in laboratory-evolved microbes from next-generation sequencing data using breseq. Methods Mol. Biol. Clifton NJ 1151, 165–188 (2014).

59. Langmead, B. & Salzberg, S. L. Fast gapped-read alignment with Bowtie 2. Nat. Methods 9, 357–359 (2012).

