## Supplemental data for "Differential evolution of cooperative traits in aggregative multicellular bacterium *Myxococcus xanthus* driven by varied population bottleneck sizes"

**Social traits of *M. xanthus***

**Sporulation**: Sporulation of *M. xanthus* is triggered by starvation and results in the formation of spore filled fruiting bodies [1]. This process requires multiple contact independent signalling mechanisms such as quorum sensing [2,3], and death of a major fraction of cells die presumably to provide nutrients to the small fraction of cells that successfully develop into spores [4]. This process thus is susceptible to the evolution non-cooperating individuals as was demonstrated previously [5].

**Predation:** Predation by *M. xanthus* is mediated by both contact dependent and independent antimicrobial mechanisms [6]. Contact dependent killing mechanisms rely on the expression and use of secretion systems [7]. Whereas contact independent mechanisms are diverse and include production of antibiotics, toxins, antimicrobial peptides, digestive enzymes and secondary metabolites [8]. Moreover, prey cells are digested extracellularly and hence in addition to the antimicrobial substances that kill the prey, nutrients from the dead prey cells serve as public goods. It is easy to see how such a predatory strategy will rely on high densities of the predator and should therefore show density dependence. Recent reports demonstrate that indeed predation by *M. xanthus* is density dependent [9]. Each of the antimicrobial strategy listed above is costly to synthesize / express, thus the cells that do not produce antimicrobial substances listed above stand to gain dipropionate benefit. Taken together, *M. xanthus* predation is driven by cooperative / synergistic interactions between individuals that is susceptible to exploitation.

**Germination:** Recent evidence suggests that *M. xanthus* germination efficiency increases as a function of density of *M. xanthus* population [10] also this process is pervasive to cheating behaviours by non-cooperating cells. Cheating in this case might be driven by the variants that disproportionately take advantage of diffused public goods.

**Growth:** Similar to previous report [11]**,** we show that the growth of *M. xanthus* (swarming on agar surface) is density dependent. Casitone is a complex media, and digesting extracellularly available autoclaved casitone media involves secretion of digestive enzymes. Thus the extracellular antimicrobial molecules as well as the digested prey are public goods that are available to the producers as well as the non-producers.

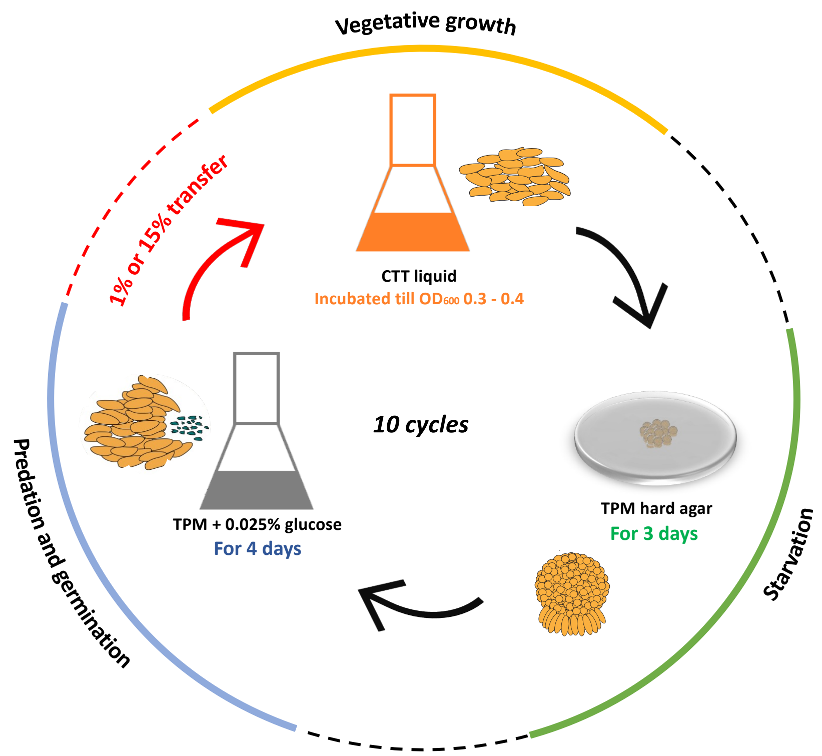


**Supplementary figure 1: Outline of the life cycle experimental evolution of *M. xanthus* with either 1 % (stringent) or 15 % (relaxed) population bottlenecks.** Four different colonies of *M. xanthus* (GV1) were used to establish four parallel evolving lines. One generation of complex lifecycle with multiple social traits involved growth of *M. xanthus* populations in nutrient rich CTT liquid (with gentamycin) medium till O.D. $_{600\ nm}$ reached 0.3 - 0.4. These cultures were then spotted on starvation TPM hard agar (1.5 % agar) plate for sporulation and fruiting body development. Next, only the spores (and not vegetative cells that failed to sporulate) were harvested by first incubating the *M. xanthus* populations at 50 °C before they were transferred on the lawn of *E. coli* for germination and predation. After incubation for 4 days on *E. coli* lawns, populations were harvested by adding 4 mL TPM buffer, shaking at 200 rpm and either (0.04 mL) 1 %, or (0.6 mL) 15 % of harvested populations were transferred to fresh CTT liquid media with gentamycin *(M. xanthus* is naturally resistant to gentamycin whereas *E. coli* is sensitive to it). This selection regimen was repeated for 10 cycles.

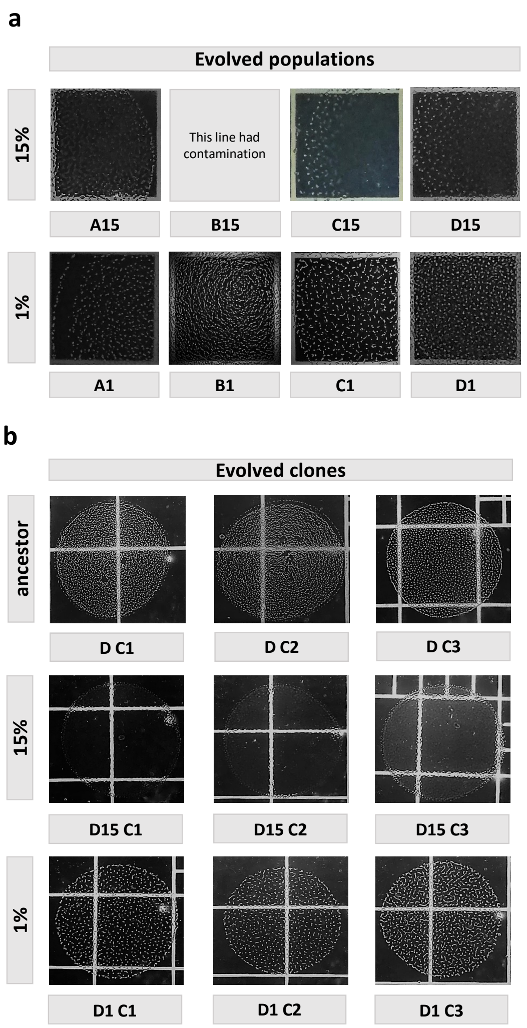


**Supplementary figure 2: Fruiting body spots of the stringent (1 %) and relaxed (15 %) populations and clones are morphologically different from each other and the ancestor.**

Representative images of the TPM hard agar (1.5 % agar) plates post three days of incubation of 100 µL of *M. xanthus* $10^9$ cells / mL density cultures are shown. Small white dot represent individual fruiting body after 3 days of incubation. (a) Populations from relaxed regimen (15 %) had fewer fruiting bodies, whereas populations from stringent bottleneck regimen (1 %) exhibited proficient fruiting body formation (small white dots). (b) Similar to population level observation, the clones isolated from D15 line (representative line from relaxed regime) was less efficient at fruiting body formation, whereas the clones from D1 (representative line from stringent regime) exhibited proficient fruiting body development.

| Reference genome | Position | Mutation | Gene | Clone | Evolution condition | Type | Effect | Codon change | Amino acid change | MXAN locus tag | Additional information | Reference |
|---|---|---|---|---|---|---|---|---|---|---|---|---|
| DK1622 | 35,62,446 | T -> C | Intergenic | • D1C1<br>• D1C2<br>• D1C3 | 1% | SNP | - | - | - | - | - | - |
| DK1622 | 60,70,872 | (T)6 –> (T)7 | transposase orfB, IS3 family | • D1C3 | 1% | SNP | Frameshift | TTC -> TTT | K96K | MXAN_4850 | - | - |
| DK1622 | 61,30,945 | (A)8 –> (A)9 | sigma 54-interacting transcriptional regulator | • D1C1<br>• D1C2<br>• D1C3 | 1% | SNP | Frameshift | AAC -> AAA | N444K | MXAN_4899 | Direct role in secondary metabolite production : **potential role in predation** | Volz C. et.al., ChemBiol (2012) doi: 10.1016/j.chembiol.2012.09.010 |
| DK1622 | 83,45,208 | C -> A | LysR family transcriptional regulator | • D1C2<br>• D1C3 | 15% | SNP | Missense | CGC -> AGC | R145S | MXAN_6792 | Uncharacterized protein | Goldman B. S. et.al., PNAS (2006) doi: 10.1073/pnas.0607335103. |
| DK1622 | 60,70,872 | (T)6 –> (T)7 | transposase orfB, IS3 family | • D15C2 | 15% | SNP | Frameshift | TTC -> TTT | K96K | MXAN_4850 | - | - |
| DK1622 | 12,73,407 | C -> T | DNA binding response regulator | • D15C1<br>• D15C2<br>• D15C3 | 15% | SNP | Missense | ACG -AUG | T75M | MXAN_1093 | Direct role in fruiting body development : **potential reason for decreased sporulation** | Peters T. et.al., Mol. Microbio. (2012) doi: 10.1111/j.1365-h2958.2012.08015.x. |

**Supplementary Table 1: Distinct mutations in stringent and relaxed bottleneck evolution regimes.** Whole genome sequencing was performed on representative clones from the stringent (1 %) and relaxed (15 %) regimens. All sequenced clones from stringent regimen had a frameshift mutation in one of the sigma 54 interacting transcriptional regulators (MXAN_4899). All sequenced clones from relaxed regimen had a missense mutation in one of the DNA binding response regulator genes (MXAN_1093).

| Clone | Sporulation efficiency (%) | Confidence interval (95%) |
|---|---|---|
| **Ancestor (clone 1, DC1)** | **1.283** | **0.005** |
| **Ancestor (clone 2, DC1)** | **1.061** | **0.004** |
| **Ancestor (clone 3, DC1)** | **1.197** | **0.006** |
| **15 % (clone 1, D15C1)** | **0.002** | **0.00003** |
| **15 % (clone 2, D15C2)** | **0.004** | **0.00005** |
| **15 % (clone 3, D15C3)** | **0.003** | **0.00005** |
| **1 % (clone 1, D1C1)** | **0.893** | **0.004** |
| **1 % (clone 2, D1C2)** | **1.616** | **0.008** |
| **1 % (clone 3, D1C3)** | **1.140** | **0.005** |

**Supplementary Table 2: Clones from relaxed bottleneck regime show drastically reduced sporulation efficiency compared to the clones from both stringent and ancestor regime.**

The table represents the sporulation efficiencies of three clones of stringent (1 %), relaxed (15 %) and ancestor regime. The sporulation efficiencies are calculated as the percentage of spores formed when 100 µL of 5 x $10^9$ cells / mL was allowed to sporulate on TPM hard agar (1.5 % agar) plate.

### References


1. Reichenbach, H. & Dworkin, M. The order Myxobacterales. in the prokaryotes: a handbook on habitats, isolation, and identification of bacteria (eds. Starr, M. P., Stolp, H., Trüper, H. G., Balows, A. & Schlegel, H. G.) 328–355 (Springer, 1981). doi:10.1007/978-3-662-13187-9_20.
2. Kaplan, H. B. & Plamann, L. A *Myxococcus xanthus* cell density-sensing system required for multicellular development. *FEMS Microbiol. Lett.* **139**, 89–95 (1996).
3. Shimkets, L. J. Intercellular signaling during fruiting-body development of *Myxococcus xanthus*. *Annu. Rev. Microbiol.* **53**, 525–549 (1999).
4. Wireman, J. W. & Dworkin, M. Developmentally induced autolysis during fruiting body formation by *Myxococcus xanthus*. *J. Bacteriol.* **129**, 798–802 (1977).
5. Velicer, G. J., Kroos, L. & Lenski, R. E. Developmental cheating in the social bacterium *Myxococcus xanthus*. *Nature* **404**, 598–601 (2000).
6. Berleman, J. E. & Kirby, J. R. Deciphering the hunting strategy of a bacterial wolfpack. *FEMS Microbiol. Rev.* **33**, 942–957 (2009).
7. Seef, S. *et al.* A Tad-like apparatus is required for contact-dependent prey killing in predatory social bacteria. *eLife* **10**, e72409 (2021).
8. Goldman, B. S. *et al.* Evolution of sensory complexity recorded in a myxobacterial genome. *Proc. Natl. Acad. Sci. U. S. A.* **103**, 15200–15205 (2006).
9. Muñoz-Dorado, J., Marcos-Torres, F. J., García-Bravo, E., Moraleda-Muñoz, A. & Pérez, J. *Myxobacteria*: Moving, Killing, Feeding, and Surviving Together. *Front. Microbiol.* **7**, 781 (2016).
10. Pande, S., Pérez Escriva, P., Yu, Y.-T. N., Sauer, U. & Velicer, G. J. Cooperation and cheating among germinating spores. *Curr. Biol* **30**, 4745-4752.e4 (2020).
11. Rosenberg, E., Keller, K. H. & Dworkin, M. Cell density-dependent growth of *Myxococcus xanthus* on casein. *J. Bacteriol.* **129**, 770–777 (1977).
